# Combining Top-Down and Bottom-Up: An Open Microfluidic Microtumor Model for Investigating Tumor Cell-ECM Interaction and Anti-Metastasis

**DOI:** 10.1101/2024.03.19.585776

**Authors:** Chao Li, Jiayi Li, Zach Argall-Knapp, Nathan W. Hendrikse, Mehtab A. Farooqui, Bella Raykowski, Anna King, Serratt Nong

**Affiliations:** Carbone Cancer Center, University of Wisconsin-Madison, 53792 Madison, WI (USA); Department of Biomedical Engineering, University of Wisconsin-Madison, 53706 Madison, WI (USA); College of Osteopathic Medicine, Liberty University, 24502 Lynchburg, VA (USA); Department of Biochemistry, University of Wisconsin-Madison, 53706 Madison, WI (USA); Department of Integrative Biology, University of Wisconsin-Madison, 53706 Madison, WI (USA)

**Keywords:** Tumor Microenvironment, Open Microfluidics, ECM Alignment&Remodeling, Cell Migration, Anti-Metastasis

## Abstract

Using a combined top-down (i.e., operator-directed) and bottom-up (i.e., cell-directed) strategy, we present an Under-oil Open Microfluidic System (UOMS)-based microtumor model for investigating tumor cell migration and anti-metastasis drug test. Compared to the mainstream closed microfluidics-based microtumor models, the UOMS microtumor model features: i) micrometer-scale lateral resolution of surface patterning with open microfluidic design for flexible spatiotemporal sample manipulation (i.e., top-down); ii) self-organized extracellular matrix (ECM) structures and tumor cell-ECM spontaneous remodeling (i.e., bottom-up); and iii) free physical access to the samples on a device with minimized system disturbance. The UOMS microtumor model is used to test an anti-metastasis drug (incyclinide, aka CMT-3) with a triple negative breast cancer cell line (MDA-MB-231). The *in vitro* results show a suppression of tumor cell migration and ECM remodeling echoing with the *in vivo* mice metastasis results.

## Introduction

For over a decade, bioengineers have been dedicated to constructing microtumor models *in vitro* that recapitulate the *in vivo* tumor microenvironment (TME).^[1]^ *In vitro* microtumor models provide a venue for using human cell lines or primary patient samples (distinct from animal models with a non-human environment)^[2]^ to perform pathogenesis studies, diagnosis/prognosis, and drug/treatment screening.^[3]^ In light of the value in both fundamental research and translational application, recently *in vitro* microtumor models got further developed to better mimic the *in vivo* physiopathological environments, witnessing the transition from the traditional two-dimensional (2D) cell culture [i.e., cells directly cultured on an artificial substrate such as polypropylene (PS) or glass]^[4]^ to three-dimensional (3D) microtissues (e.g., spheroids,^[5]^ organoids^[6]^) supported by ECM^[7]^ and microphysiological systems^[8]^ with advanced structures [e.g., lumen,^[9,10]^ microvascular networks (MVNs)^[11]^] and mass transport (e.g., vital gasses,^[12]^ soluble factors^[13]^) controls.

So far various strategies have been developed to establish a TME *in vitro*, falling into two categories regarding the design principles - top down and bottom up. For top down, the microenvironment is defined and controlled by the operators, i.e., operator-directed.^[9,10,14]^ By contrast and in a bottom up approach, the microenvironment is self-organized and regulated by cells as seen *in vivo*, i.e., cell-directed.^[15]^ The top down approaches^[16]^ allow geometrically precise dimensions and structures, inherently pursuing an operator-directed, stable microenvironment (e.g., spatiotemporal sample organization, nutrient level, vital gas availability and kinetics, fluid dynamics) with the environment parameters largely monitored and controlled at the global level (e.g., temperature, oxygen, carbon dioxide in standard cell culture incubators, flow rate control by syringe pumps). However, the real TMEs in the body are a highly dynamic and heterogeneous environment featuring aberrant structures (e.g., leaky microvasculatures, ECM remodeling) and mass transport dynamics (e.g., nutrients, oxygen, metabolic wastes) defined by the cells locally in the micro niche. The bottom up approaches^[17]^ give the driver’s seat back to the cells in a micro niche and therefore better mimic the *in vivo* processes.

A trend of using top down and bottom up combinedly to establish a TME *in vitro* has seen a fast growth in the past five years. In the combined top down and bottom up approaches, the operators define the system environment and structures at macrotissue scales (typically >100 μm) and in the meanwhile, cells are allowed to autonomously generate and regulate their own micro niche and fine structures (e.g., stem cell crypts,^[18]^ MVNs^[19]^) at microtissue and single-cell scales (typically <100 μm). In this regime, the advantages from top down (i.e., macrotissue-level precision and controllability) and bottom up (i.e., single-cell-level structures and autonomously regulated microenvironment dynamics) get integrated and synergized in one system.

In the combined top down and bottom up approaches, microfluidic systems especially closed-chamber/channel microfluidic systems (i.e., fluid/cells confined in a closed space by solid barriers) dominate due to the well-established fabrication methods and fluid controls.^[20]^ While successful in many aspects, the closed-chamber/channel microfluidic systems are inherently limited by their closed system configuration that is misaligned with the open system standard (e.g., pipette, glass slide, chambered coverglass, Petri dish, microtiter plate) systematically adopted in biology and biomedicine. Specifically, there is no easy and non-destructive physical access to the samples on a closed device. The closed system configuration makes real-time spatiotemporal sample organization on a device (e.g., for sample/drug loading, manipulation, and collection) challenging and disruptive to the established TME.

Here, we leverage a newly introduced sub-branch in open microfluidics known as Exclusive Liquid Repellency (ELR)-empowered Under-oil Open Microfluidic System (UOMS)^[12,21–28]^ to establish an open microfluidic microtumor model with combined top down and bottom up. The ELR physics allows inherent (i.e., surface-texture and surfactant independent) and absolute liquid repellency (with Young’s contact angle *θ* = 180o),^[21,23]^ which lays the foundation for the top down fabrication of open microfluidic channels in UOMS with micrometer-scale spatial resolution and lossless sample processing. Furthermore, ECM materials (e.g., collagen Type I) can be introduced into the microchannels via a unique microfluidic sample distribution method named “under-oil sweep distribution”^[24,25]^ and self-organize into a structure bottom up with different levels of microfiber alignment from random to aligned via a Laplace-pressure-driven local lateral flow.^[25]^ Tumor cells seeded through the oil overlay onto the under-oil microchannels interact with the ECM layer, further remodeling the ECM structure bottom up which in turn influences cell morphology and migration. Thanks to the oil-media barrier, the UOMS microtumor model allows real-time, free physical access to the system with minimized system disturbance (e.g., media loss via evaporation, airborne contamination, irreversible and destructive intervention). In this work, we present the governing physics, design principle, and self-organization mechanism of the under-oil TME. As a proof of principle, we test the UOMS microtumor model with a repurposed anti-metastasis drug [incyclinide, a matrix metalloproteinase (MMP) inhibitor that went through several clinical trials for brain tumors, soft tissue tumors, and metastatic cancer treatment], showing its successful capture of the drug efficacy consistent with the *in vivo* (mice model) results.

## Results and Discussion

Microfluidic cell culture has many advantages over the conventional bulk-scale cell culture (e.g., Petri dish, microtiter plate) including microstructure development and mass transport control for better mimicking the dynamic and heterogeneous cellular microenvironments as seen *in vivo*.^[20]^ In microfluidics, closed-chamber/channel microfluidic systems dominate due to the historical reason and intuitive fluid controls (i.e., fluid in a closed space defined by solid barriers). However, in biology and biomedicine, open systems (such as pipette, glass slide, chambered coverglass, Petri dish, microtiter plate) are systematically adopted. To better align with the open system standard, open microfluidics [i.e., fluid in an open space with at least one solid barrier removed, exposing the fluid to the ambient air (single-liquid-phase microfluidics) or a secondary fluid (multi-liquid-phase microfluidics) for free physical access to the system] were introduced.^[29]^ While promising in biological and biomedical applications, a broad adoption of open microfluidic systems had been stymied (before 2015) due to the limited spatial resolution (largely limited to millimeter scale), system stability (e.g., media loss via evaporation, airborne contamination), fluid controls (e.g., low and limited flow rate range, lack of on-demand flow switch) compared to closed-chamber/channel microfluidics.

### Top down - Double-ELR open microfluidic surface patterning

Recently, we introduced ELR (i.e., Exclusive Liquid Repellency)^[21]^ - an extreme wettability where a liquid is inherently and absolutely repelled on a surface (Figure 1) - to open microfluidics, initiating a new sub-branch known as ELR-empowered UOMS. Enabled by ELR^[21]^ and Double-ELR (i.e., under-oil water ELR + under-water oil ELR on a chemically patterned surface),^[24]^ the ELR-empowered UOMS^[25]^ achieved a series of advanced functions in open microfluidics including: i) free physical access to the system with minimized system disturbance, ii) inherent antifouling and lossless sample processing, iii) micrometer-scale spatial resolution of open microchannels, iv) open-fluid single-cell trapping, v) improved flow rate range (e.g., comparable to blood flow in the circulatory systems), and vi) on-demand reversible open-fluid valves. These advances not only make ELR-empowered UOMS comparable to closed-chamber/channel microfluidics in functionality but also enables new applications in biology and biomedicine.^[26,27]^

**Figure 1.**
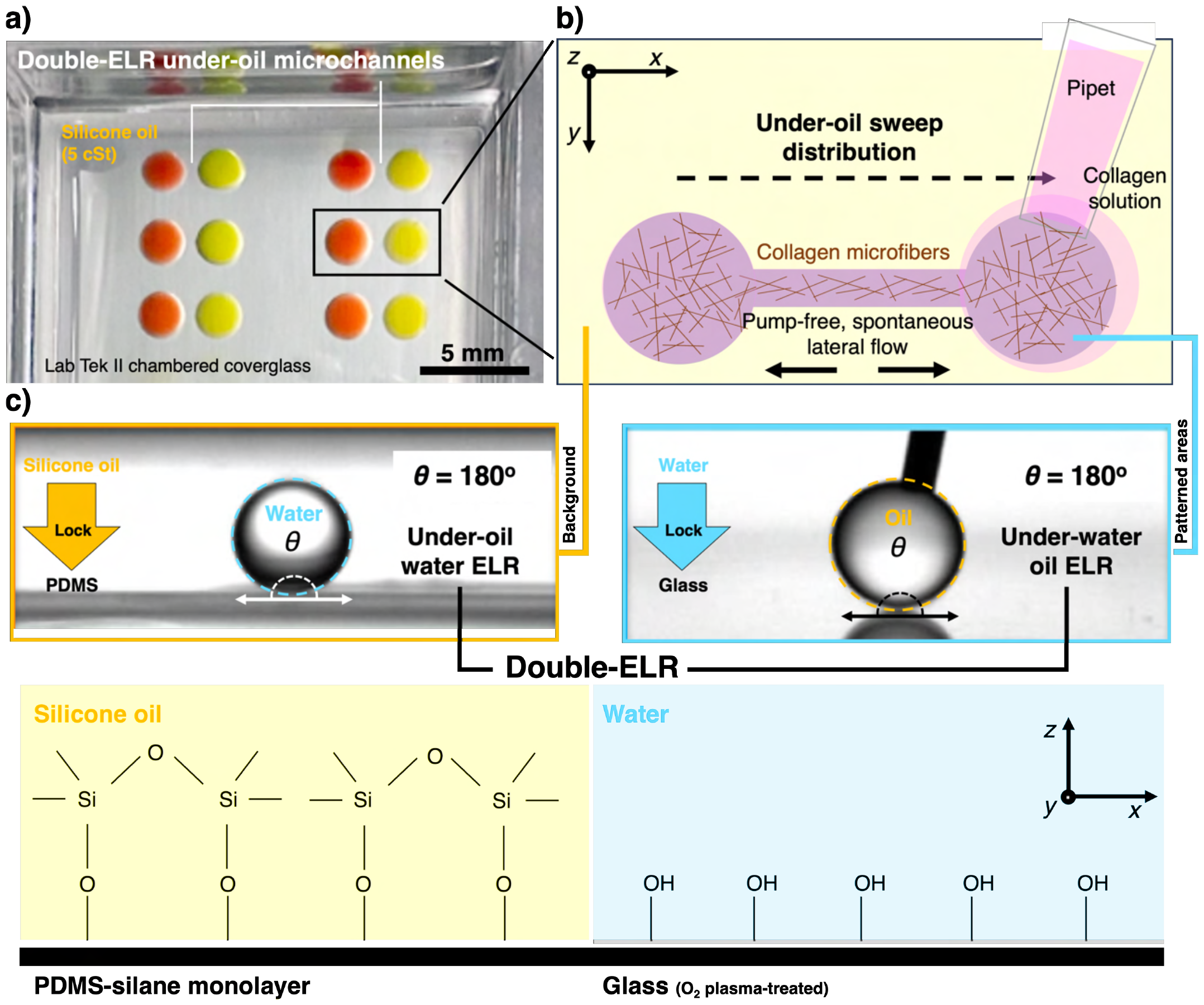
ELR physics and under-oil sweep distribution for preparing under-oil microchannels with ECM coating. (a) A camera photo of the Double-ELR [i.e., under-oil water ELR (background) + under-water oil ELR (patterned areas)] under-oil microchannel device [Lab Tek II chambered coverglass overlaid with silicone oil (5 cSt)]. The microchannel dimensions are written in “2-2-*L*0.5-*W*0.1” for 2 mm of the spot diameter (*D*spot), 0.5 mm of the channel length (*L*), and 0.1 mm of the channel width (*W*). The liquid used here is RPMI+10% FBS with food dye (red and yellow) for visualization. (b) A schematic (top view, *xy* plane) showing the under-oil sweep operation. Collagen solution (see Experimental Section) is dispensed onto the patterned areas by sweeping a hanging drop across the surface with pipet at room temperature (RT, ∼21 °C). Collagen microfibers form and self-assemble over time during polymerization (RT/1 h) and get aligned in the microchannel driven by the spontaneous lateral flow triggered by the local Laplace pressure differential between the channel and the spots. (c) Physics and surface chemistry of Double-ELR (side view, *xz* plane). (Top) The goniometer images showing the 180o Young’s contact angle (*θ*). The double-headed arrows indicate the balanced interfacial tensions (*γ*) of the solid-oil-water three phases. The dashed-line circles highlight the spherical profile of the droplets (3 μL). Oil and water (or cell culture media) are absolutely “locked” on their preferred surface (i.e., PDMS for oil, glass for water). (Bottom) The surface chemistry of each surface. The background is a monolayer of PDMS-silane molecules covalently bonded to the surface. The patterned areas are O_2_ plasma-treated glass surface.

Enabled by Double-ELR, a selected pattern (e.g., two circular spots connected by a straight channel adopted in this study) can be readily prepared on a surface [e.g., glass, polystyrene (PS)] by under-oil sweep distribution (Figure 1a and 1b, Experimental Section). A hanging drop of the target media (e.g., collagen solution) at the end of a pipet tip is dragged across the Double-ELR chemically patterned surface (Figure 1c). The aqueous media gets spontaneously and robustly dispensed onto the patterned area, leaving the background (i.e., the unpatterned area) absolutely protected from being fouled. The sweep operation is performed under oil, minimizing volume loss via evaporation and airborne contamination. The spatial resolution of the surface patterns is defined by the polydimethylsiloxane (PDMS) stamps (or masks) prepared with standard photolithography (Experimental Section).

### Bottom up - Self-organization of ECM microstructure driven by spontaneous lateral flow

Right after under-oil sweep, all the liquid takes a near constant height-to-width (*H*/*W*) ratio of 1/13.^[25]^ Due to the Laplace pressure (Δ*P*) differential between the microchannel and the inlet/outlet spots (Δ*P*_microchannel_ > Δ*P*_spot_) (Table S1):

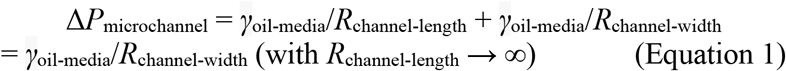

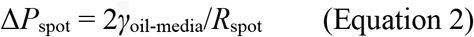

where *γ*_oil-media_ is the interfacial tension at the oil-media interface [41.8 mN/m for silicone oil (5 cSt)-water interface];[^21]^ *R*_channel-length_ and *R*_channel-width_ are the radii of curvature of the microchannel in the channel length and width direction, respectively; *R*_spot_ is the radius of curvature of the inlet/outlet spots), a lateral flow is spontaneously generated, pumping partial of the volume in the microchannel (*V*_pumping_) into the connecting spots until the system reaches pressure equilibrium (Figure 2a). The lateral flow can be qualitatively described by Poiseuille’s law as:

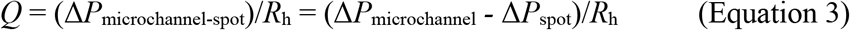

where Δ*P*_microchannel-spot_ is the Laplace pressure differential between the microchannel and the spots; *R*_h_ is the hydrodynamic resistance of the under-oil micro-channel. Here, *R*_h_ is calculated based on the following assumptions: i) the fluid is incompressible, Newtonian, and in laminar flow with a non-slippery boundary,^[30]^ and ii) the microchannel takes a constant, rectangular cross-section.

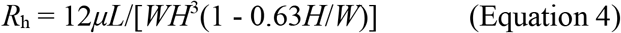

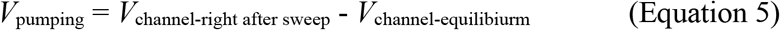

where *μ* is the viscosity of the liquid (3.84 Pa·s for collagen molecules)^[31]^. *L* is channel length, *W* is channel width, and *H* is channel height (1/13*W* right after sweep). The instant linear velocity of the lateral flow (*v*) can be known as:

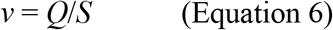

where *S* is the cross-sectional area of the microchannel perpendicular to the channel length direction at a time point (Table S1). It is worth noting that the spontaneous lateral flow generated by Laplace pressure differential between the microchannel and the spots decreases over time until Δ*P*_microchannel-spot_ = 0 Pa when the system reaches pressure equilibrium. The time (*t*) of lateral flow to reach equilibrium can be estimated as (by assuming a steady flow):

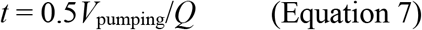

**Figure 2.**
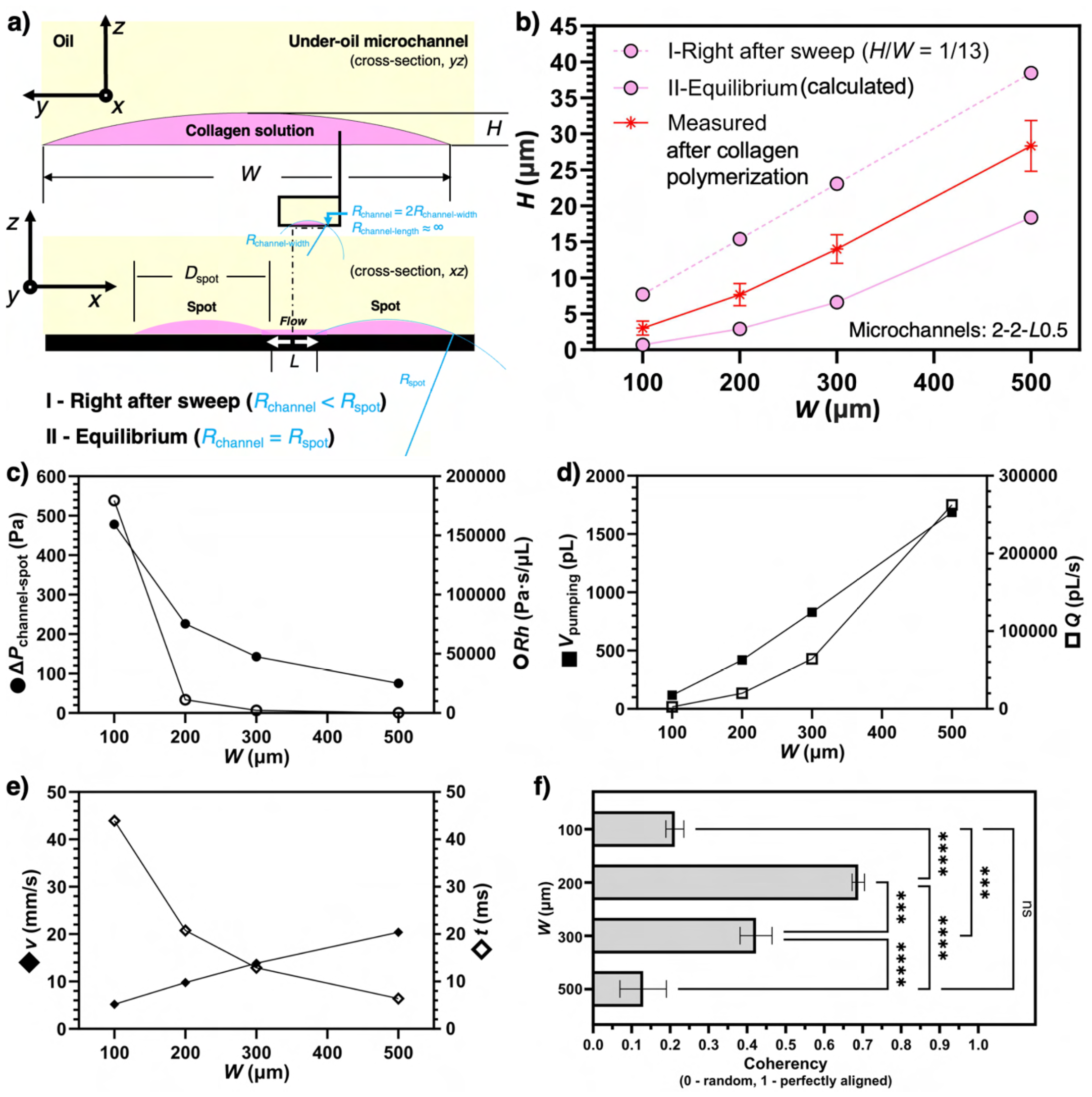
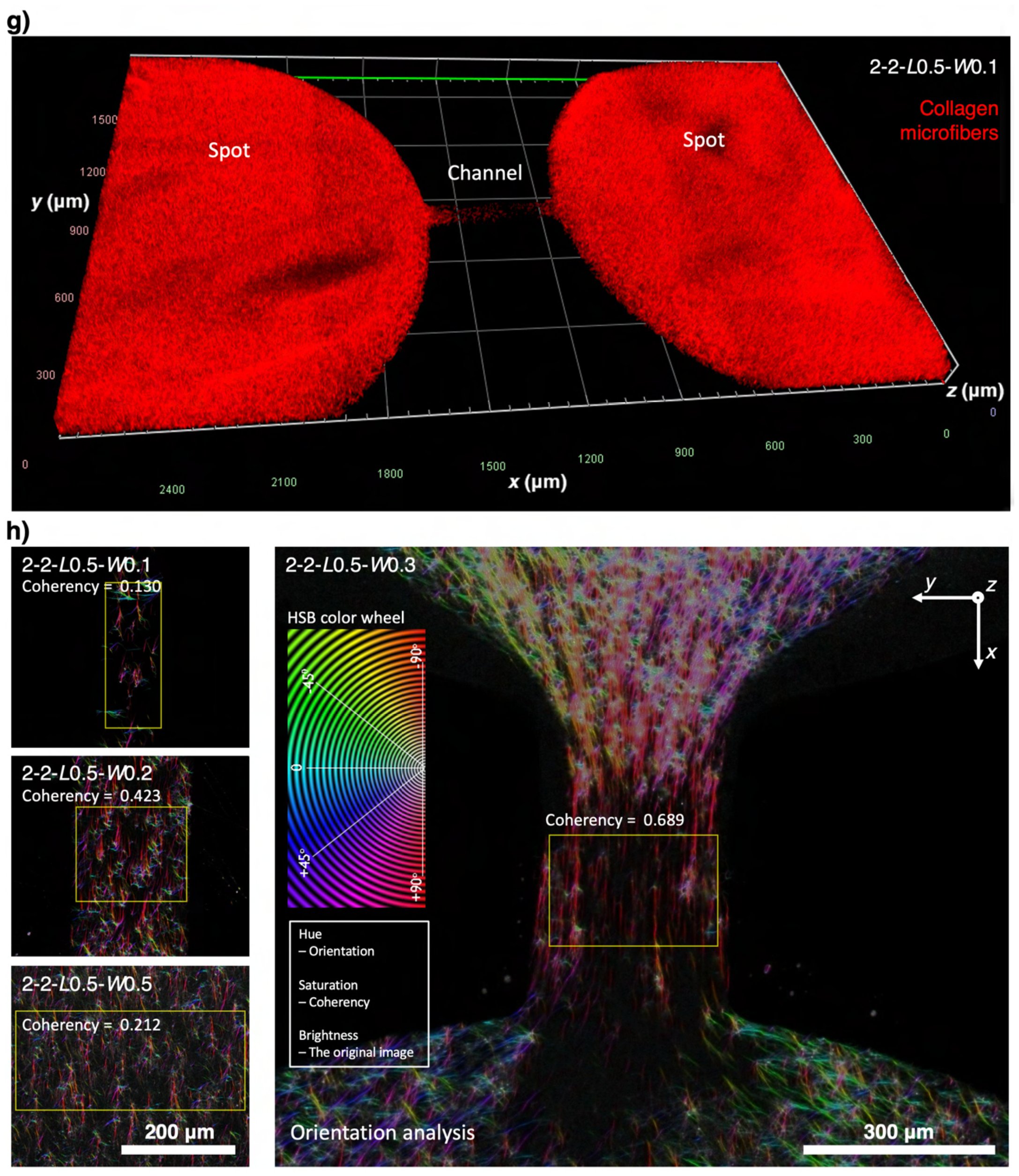
Self-organization of ECM (collagen I) microfibers in the under-oil microchannels driven by spontaneous lateral flow. (a) Fluid dynamics during and after under-oil sweep distribution. After sweep, the volume in the channel gets partially pumped into the spots until the system reaches Laplace pressure equilibrium. (b) Channel height as a function of channel width for 2-2-*L*0.5. The thickness of the ECM layer defined by the channel width was measured on Zeiss Apotome 3 (Figure S1). (c), (d), and (e) Fluid dynamic parameters as a function of channel width with a given channel length and spot size. (f) ECM microfiber alignment results showing the coherency of the alignment as a function of channel width. (g) A representative global view of 3D microscopic image (Zeiss Apotome 3) of the under-oil microchannel with ECM (red) coating. (h) Color survey of collagen microfiber orientation analysis (see Experimental Section).[^48]^ (Inset) The hue-saturation-brightness (HSB) color wheel of the orientation analysis [hue - orientation, saturation - coherency (0 - random, 1 - perfectly aligned), brightness - original image]. The yellow boxes indicate the ROIs in the orientation analysis. Error bars are mean ± S.D. from ≥3 replicates. ^*^*P* ≤ 0.05, ^*^\**P* ≤ 0.01, ^*^\*\**P* ≤ 0.001, and ^*^\*\*\**P* ≤ 0.0001. “ns” represents “not significant”.

Collagen solution starts to polymerize and self-assemble into nanofibrils and microfibers after under-oil sweep distribution at room temperature (RT, ∼21 °C). The measured thickness of the collagen layer falls between the theoretically calculated initial media layer thickness and the equilibrium media layer thickness (Figure 2b, Figure S1). The greater measured thickness compared to the equilibrium thickness can be attributed to the low mobility of the collagen colloidal solution (*μ*_collagen molecule_ = 3.84 Pa·s; *μ*_collagen fibril_ = 0.09-1.63 GPa·s) compared to water (*μ*_water_ = 1 mPa·s). Once the collagen solution is mostly polymerized, the lateral flow is gone and the thickness of the collagen layer is “frozen” before it can reach the calculated equilibrium thickness.

Here, we first analyze the fluid dynamics of a set of microchannels with the same channel length (*L* = 0.5 mm) and varying channel width (*W* = 0.1, 0.2, 0.3, and 0.5 mm) (Figure 2c to 2h). The analytical results showed that Δ*P*_microchannel-spot_ and *R*_h_ (Figure 2c) increase with *R*_h_ much faster than Δ*P*_microchannel-spot_ as channel width decreases, resulting in a decrease of *Q* (Figure 2d) quickly. From after under-oil sweep distribution to equilibrium, *V*_pumping_ (Figure 2d) gets spontaneously pumped into the two spots. The direct driving force of collagen microfiber alignment in the microchannel is the shear force generated by the spontaneous lateral flow. The intensity of shear force can be quantitatively reflected by *v* (Figure 2e). The duration of shear force is described by *t* (Figure 2e). By leveraging the spontaneous lateral flow, different levels of collagen microfiber alignment from random to aligned can be achieved by simply adjusting the channel dimensions (e.g., channel width in this section) (Figure 2f to 2h). For a given channel length (e.g., *L* = 0.5 mm), the microfiber alignment increases first and then decreases as the channel width decreases. This trend can be understood by considering *v* and *t* jointly (Figure 2e). A larger channel width leads to a higher *v* (and thus shear force), however, a shorter *t* for the microfibers to align with the flow. As shown in both the analytical and experiential results, the highest alignment is achieved with the microchannels of 2-2-*L*0.5-*W*0.3 (in mm) (Figure 2h). From *W* = 0.3 mm, both increase and decrease of channel width leads to decreased microfiber alignment due to either not enough time or not enough linear velocity of the lateral flow.

Next, we further investigate the influence of channel length on the ECM microfiber alignment in the under-oil microchannels (Figure S2). As predicted by the under-oil fluid dynamics analysis (Figure 2c to 2e) and for a given channel width (and thus the same Δ*P*_microchannel-spot_), increase of channel length will lead to a decrease of *v* but an increase of *t*. Therefore, the increase of channel length should result in an increase of microfiber alignment but eventually decrease if the channel length gets too large. This hypothesis is proved by the experimental results (Figure S2b to 2c). It is worth noting that the lateral flow generation and dynamics in UOMS are fundamentally different from the traditional closed microchannels with active pumping.^[32]^ Right after sweep, the lateral flow is driven by the “local” Laplace pressure differential rather than the global pressure drop by active pumping. A part of the volume gets pumped into the two spots, generating spontaneous later flows in two opposite directions in the channel (Figure 1b, Figure 2a). Due to the orders-of-magnitude difference in the fluid volume between the spots and the microchannels (Table S1), the lateral flow gets significantly damped after reaching the spots, leading to much weaker to no microfiber alignment effect compared to the microchannels (Figure S2d). Together, this unique under-oil fluid microenvironment allows us to get different ECM microfiber alignments by simply selecting the channel length, channel width, and spot diameter. Via an under-oil sweep or multiple layer-by-layer sweeps (Figure S3), the spontaneous lateral flow drives the ECM microfibers into a specific level of alignment during the polymerization process. This feature provides an efficient and high-throughput approach to reconstructing an ECM microenvironment that recapitulates the typical structures seen in either healthy tissues or a cancer lesion.

### Bottom up - Tumor cell/fibroblast-driven spontaneous ECM remodeling

It is known that the cells in the TME including tumor cells and fibroblasts can strongly remodel the collagen microfiber alignment, facilitating tumor cell migration and metastasis.^[33]^ In cancers, ECM remodeling and alignment often promotes tumor growth, invasion, and metastasis by providing structural support and guiding cellular movement.^[7]^ Additionally, aligned ECM can facilitate resistance to chemotherapy and immunotherapy by creating physical barriers that hinder drug penetration and immune cell infiltration.^[34]^ Understanding the relationship between ECM remodeling/alignment and tumor cell behavior has led to the development of novel treatment strategies targeting tumor-ECM interaction to enhance drug delivery and improve therapeutic outcomes, highlighting the significance of ECM in cancer progress and treatment.^[35]^

In this section, we investigate the ECM remodeling capability and capacity by tumor cells [MDA-MB-231 - a migratory breast cancer cell line, green fluorescence protein (GFP)-labeled] with and without fibroblasts [healthy human oral fibroblasts (HorF) versus cancer-associated fibroblasts (CAF), 1:1 ratio with tumor cells] using the UOMS microtumor model (Figure 3). Two seeding densities of MDA-MB-231 were tested at 5000 (i.e., 5K) cells/μL and 20K cells/μL. The MDA-MB-231 cells were added onto one of the spots (the proximal spot) of the microchannels [2-2-*L*0.4-*W*0.3 and 2-2-*L*0.5-*W*0.5 with ECM (collagen Type I, 2.5 mg/mL, red fluorescence protein (RFP)-labeled] under oil in 1 μL of media. At the low 5K seeding density, the tumor cell monoculture groups showed about 40% ECM contraction by 18 h. Interestingly, the 5K tumor cell-fibroblast co-culture groups showed noticeably higher ECM contraction in about 50% and 70% for CAF and HorF, respectively. In comparison, the high 20K seeding density groups (both monoculture and co-culture) all showed higher ECM contraction in about 55% to 65% compared to the 5K tumor cell monoculture groups. The influence of fibroblasts on ECM contraction in the high seeding density groups was not obvious. In all these tests (Figure 3a and 3b), the distal spot of the microchannels without cell seeding showed a stable ECM structure with no contraction. Therefore, we attributed the ECM contraction to the remodeling caused by the tumor cells and fibroblasts. In the conditions with a relatively low tumor cell density, the existence of fibroblasts in the TME apparently leads to more ECM contraction. In the conditions with a high tumor density, the ECM contraction is significant with and without the existence of fibroblasts.

**Figure 3.**
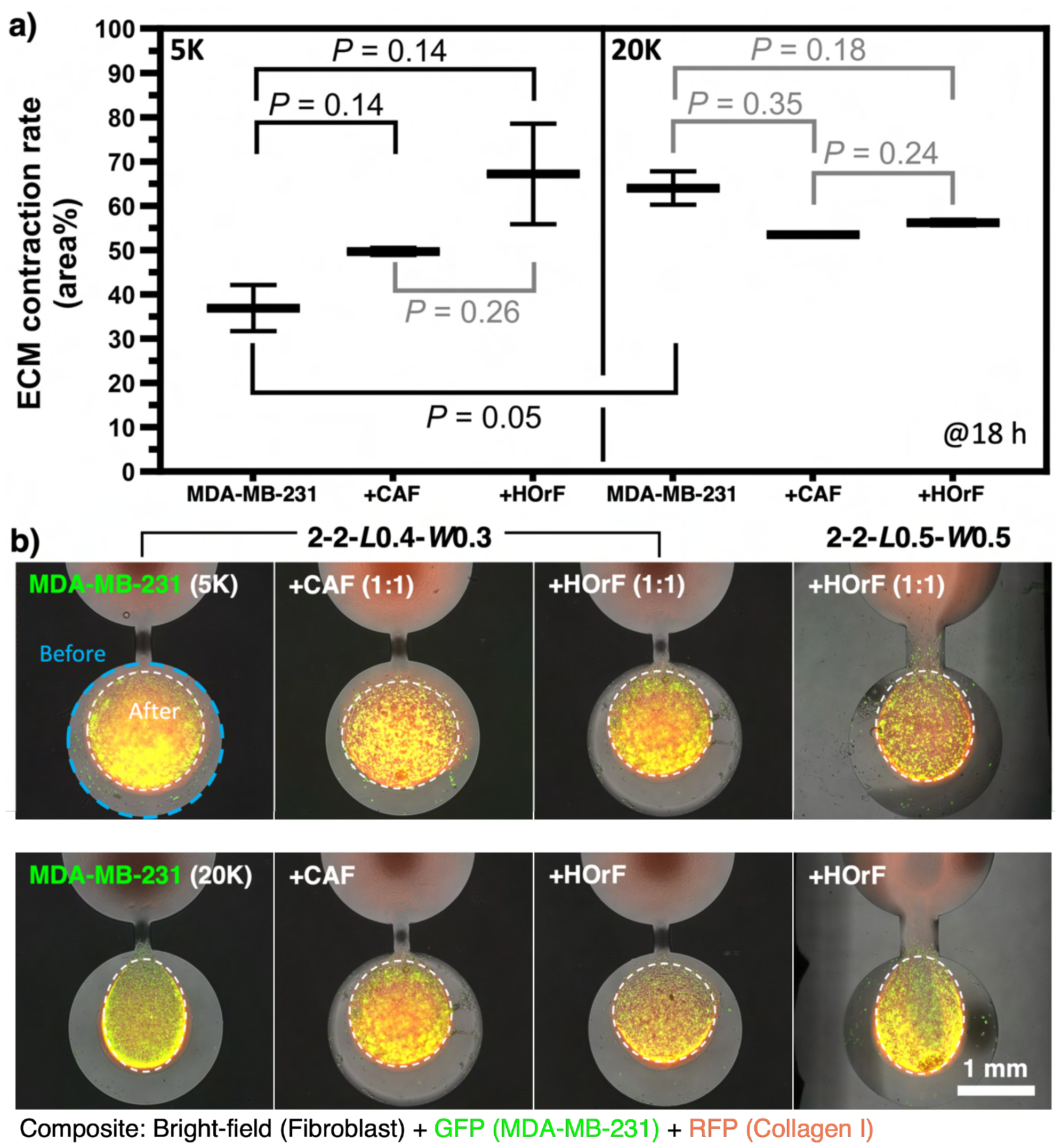
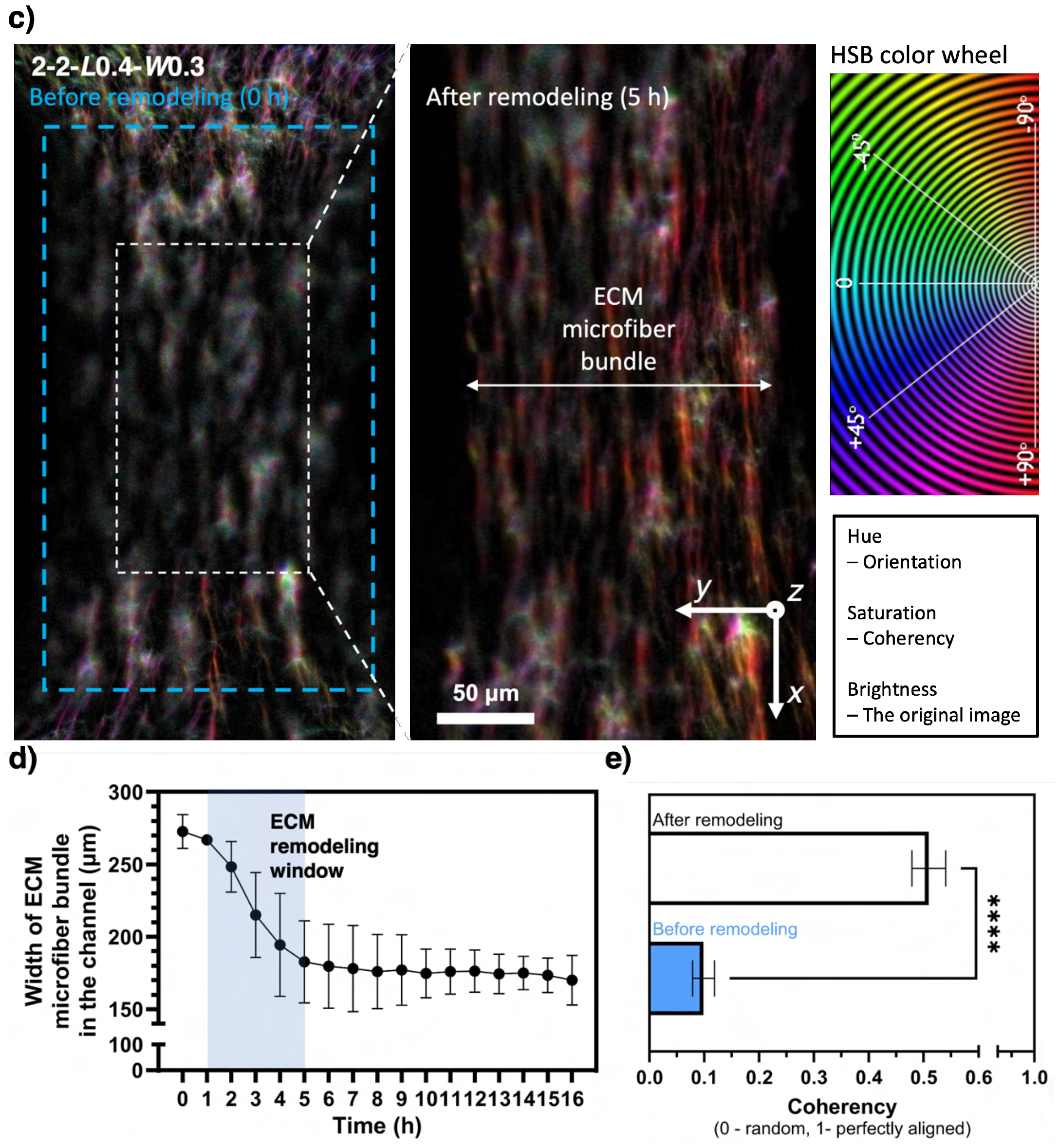
ECM contraction and remodeling driven by tumor cells with and without fibroblasts. (a) ECM contraction on the cell seeding spot (@18 h). ECM contraction rate = (area before - area after)/area before × 100%. Two seeding densities (5K versus 20K) of the tumor cells (MDA-MB-231, GFP-labeled) were tested. Fibroblasts (non-fluorescently labeled) were pre-mixed and co-cultured with tumor cells in 1:1 ratio. (b) The representative composite microscopic images corresponding to two channel dimensions - 2-2-*L*0.4-*W*0.3 and 2-2-*L*0.5-*W*0.5. The blue and white dashed-line circles indicate the ROIs before and after contraction, respectively. (c) ECM remodeling (@5 h) in the microchannel (2-2-*L*0.4-*W*0.3, Movie S1) caused by the ECM contraction on the cell seeding spot with 20K cell seeding density. The blue and white dashed-line boxes indicate the span of the microfiber bundle before and after remodeling, respectively. (d) ECM remodeling dynamics reveals an ECM remodeling window from 1-5 h (highlighted by the light blue band). (e) The coherency of the microfiber alignment in the microchannel before and after remodeling. Error bars are mean ± S.D. from ≥3 replicates. ^*^*P* ≤ 0.05, ^**^*P* ≤ 0.01, ^***^*P* ≤ 0.001, and ^****^*P* ≤ 0.0001. *P* values are specified directly on the figure if > 0.05.

Next, we further look into how the ECM contraction on the cell seeding spot may affect the ECM structure in the microchannel (Figure 3c to 3e). We chose to use the 2-2-*L*0.4-*W*0.3 microchannel to start with a low microfiber alignment in the channel (coherency <0.1) and 20K cell seeding density for a strong ECM contraction on the cell seeding spot. The results showed that at around 1 h from cell seeding the ECM contraction on the cell seeding spot started causing a visible contraction of the microfibers in the channels and quickly in about 4 h the microfibers in the channel got focused into a tight bundle (Figure S3d, Movie S1). After 5 h, the ECM remodeling in the channel reached a steady stage. Due to the strong stretching, the microfiber alignment in the channel significantly increases (coherency >0.5) (Figure 4e). If the channel width is less than 0.3 mm, the tested tumor cells at the high 20K seeding density eventually snap the microfiber bundle (see results and expended discussion in the next section, Movie S2). These ECM contraction assays demonstrated with the UOMS microtumor model can be used to evaluate the ECM remodeling capacity of primary tumor cells from patients in different cancer types and/or stages.

**Figure 4.**
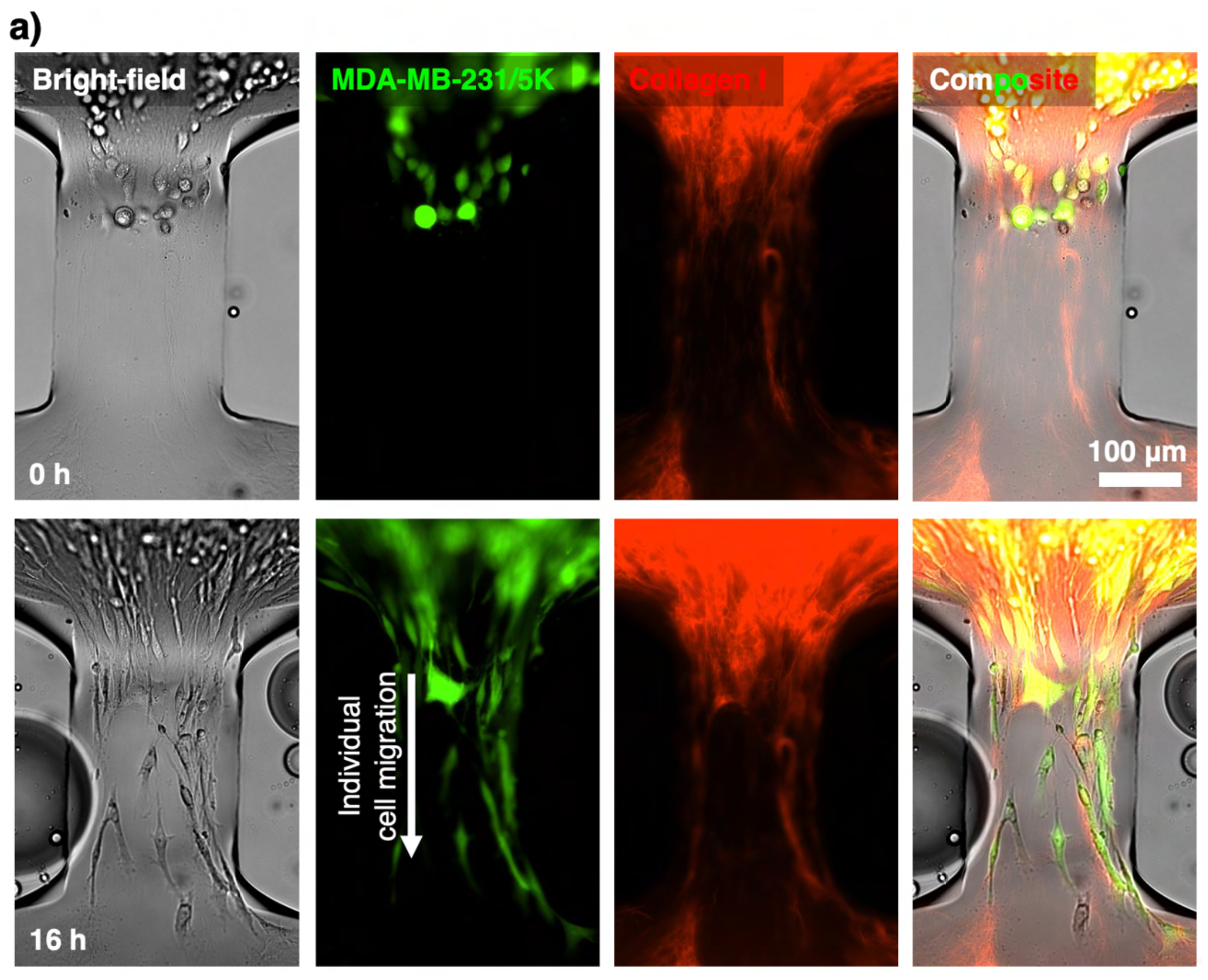
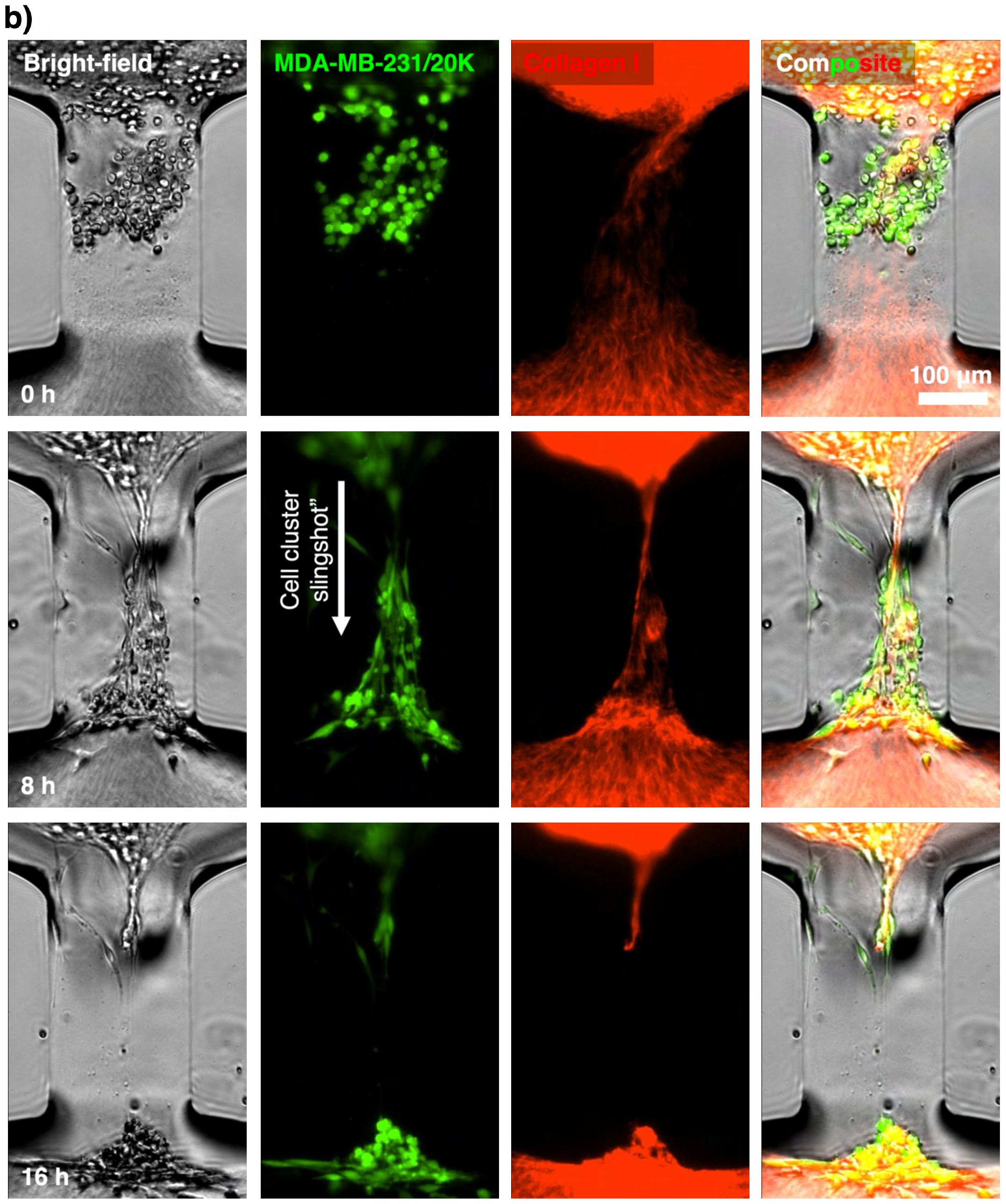
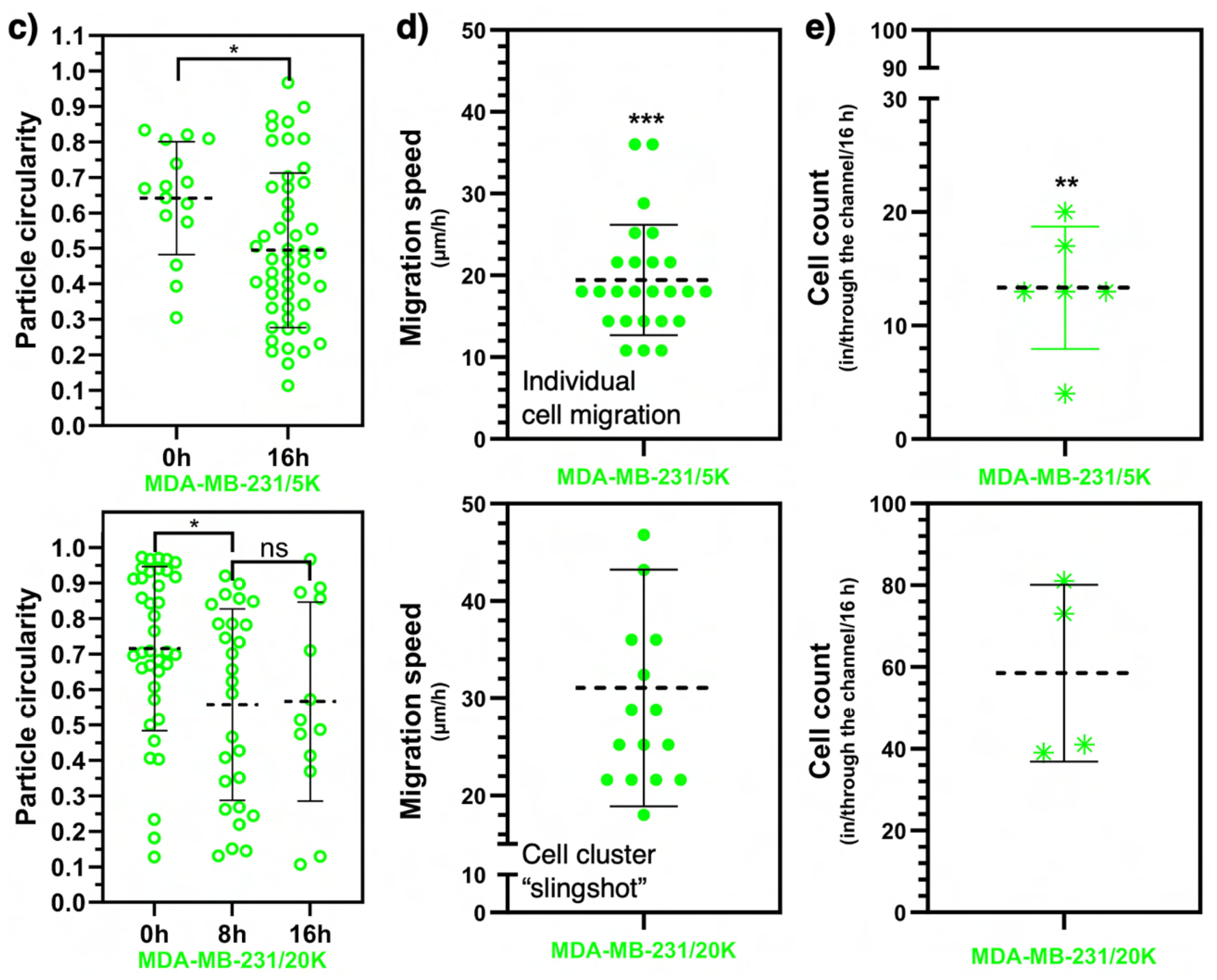
Different tumor cell migration modes captured at low and high seeding densities on the UOMS microtumor model (2-2-*L*0.3-*W*0.2). (a) Individual cell migration at low seeding density (5K) of the tumor cells (Movie S2a). (b) Cell cluster slingshot (i.e., collective migration) at high seeding density (20K) of the tumor cells (Movie S2b). (c) Cell morphology (i.e., particle circularity) analysis, (d) Migration speed analysis, and (e) Cell count analysis corresponding to the two migration modes. Error bars are mean ± S.D. from ≥3 replicates. ^*^*P* ≤ 0.05, ^**^*P* ≤ 0.01, ^***^*P* ≤ 0.001, and ^****^*P* ≤ 0.0001. “ns” represents “not significant”.

### Capture of two different modes of tumor cell migration: individual cell migration versus cell cluster slingshot

TME represents a lesioned micro niche in tissue with a complex set of stresses (e.g., low nutrients, hypoxia, high acidity) on the cells. The accumulated stresses during carcinogenesis eventually trigger epithelial-mesenchymal transition (EMT) and cancer metastasis.^[36]^ A more developed TME is expected to inflict more stresses on the cells, which could affect the process of tumor cell migration.

Here, we investigate if the UOMS microtumor model may capture any difference in tumor cell migration at different tumor cell seeding densities [i.e., low (5K cells/μL) and high (20K cells/μL) seeding densities to partially mimic the different levels of stress at the early and late tumor niche development] (Figure 4). At the low 5K seeding density, the MDA-MB-231 cells, as expected, migrated into the microchannel (2-2-*L*0.3-*W*0.2) toward the distal spot without cells. The migrating cells in the microchannels showed noticeably elongated cell morphology (>100 μm in cell length compared to about 15 μm of the cells at neutral state on the proximal spot) strictly following the aligned collagen microfibers (Figure 4a and 4c, Movie S2a). As shown in the previous section, the low seeding density causes about 40% ECM contraction during tumor cell migration in 24 h, exerting weak to little remodeling of the collagen microfibers and bundle in the microchannel.

By contrast, the MDA-MB-231 cells at the high 20K seeding density strongly contracted the ECM on the proximal spot, stretching the collage microfibers and bundle in the microchannel (2-2-*L*0.3-*W*0.2) (Figure 4b, Movie S1, Movie S2b). The strong ECM contraction heavily stretched and tightened the collagen microfiber bundle in the microchannel and finally snapped it. The tumor cells accumulated at the entrance of the microchannel got spontaneously focused into a cluster, strictly following the stretched microfiber bundle and showing a similar elongated cell morphology (Figure 4c) as observed in the 5K groups. During the snap of the microfiber bundle at the end, the whole cluster got pulled into the distal spot in a slingshot-like motion (Movie S2b). Compared to the cell migration speed of individual cells at about 20 μm/h, the cell cluster slingshot resulted in a much higher motion speed at about 32 μm/h (Figure 4d) and consequently a higher cell count reaching the distal spot (about 14 cells per microchannel for individual cell migration versus about 60 cells per microchannel for cell cluster slingshot) (Figure 4e).

As opposed to the broadly reported individual cell migration, tumor cell cluster or collective migration has been reported as an emerging tumor invasion mechanism that may reshape the understanding of cancer metastasis and anti-metastasis treatment strategies.^[37–39]^ The UOMS microtumor model successfully captures individual cell migration and a cell cluster slingshot-like motion associated with tumor cell seeding density and spontaneous tumor cell-ECM remodeling/contraction (Figure 3, Figure 4). While we preserve the caveat if similar cell cluster motion mechanisms may exist *in vivo* or not, the UOMS microtumor model demonstrates its capability to establish and investigate a variety of tumor cell-ECM microenvironments and interactions with open system standard and configurations.

### Anti-metastasis drug test on suppressing tumor cell migration and ECM remodeling

Cancer metastasis is the leading cause of mortality in cancer diseases regardless of cancer subtypes^[40]^. For example, in breast cancers, TNBC, HER-2 positive and inflammatory subtypes have poorer survival rates partially due to the higher metastatic potential.^[41]^ A drug therapy with substantial anti-metastasis efficacy may improve prognosis in patients at the early to mid stages.^[42]^

In the last section, we further demonstrate the translational capability of the UOMS microtumor model in anti-metastasis drug tests. We selected incyclinide - an MMP inhibitor - to target the ECM remodeling. Incyclinide demonstrated anti-metastatic effect in mice models inoculated with breast cancer cells (Figure 5a to 5c). Interestingly, incyclinide did not significantly decrease the primary tumor mass compared to the control group (Figure 5b, Figure S4). However, the reduction of metastasis to the lung in the treated group is statistically significant (Figure 5c). We then tested the possible anti-metastasis effect in *vitro* with the UOMS microtumor model by measuring single cell migration and ECM remodeling (Figure 5d to 5g). The microchannels we chose in this test are 2-2-*L*0.5-*W*0.1. The 0.1 mm channel width facilitates cell count and tracking in the channel (see Experimental Section) compared to thicker channels. In the UOMS microtumor model, the no-drug control groups showed strong ECM contraction on the seeding spot and high tumor cell migration into the microchannel (Figure 5e to 5g, Movie S3). By contrast, the incyclinide-treated groups showed significantly weaker ECM contraction on the seeding spot and lower tumor cell migration into the microchannel (Figure 5e to 5g, Movie S3). We acknowledge that *in vitro* microtumor models, to the best so far, only recapitulate a subset of the parameters involved in a TME *in vivo*. The anti-metastasis effect on mice might not be necessarily and completely explained by the tumor-cell ECM interactions revealed in the UOMS microtumor model, however, the UOMS microtumor model could be used as a pre-screening platform for anti-metastasis/anti-cancer drug screening before animal model tests.

**Figure 5.**
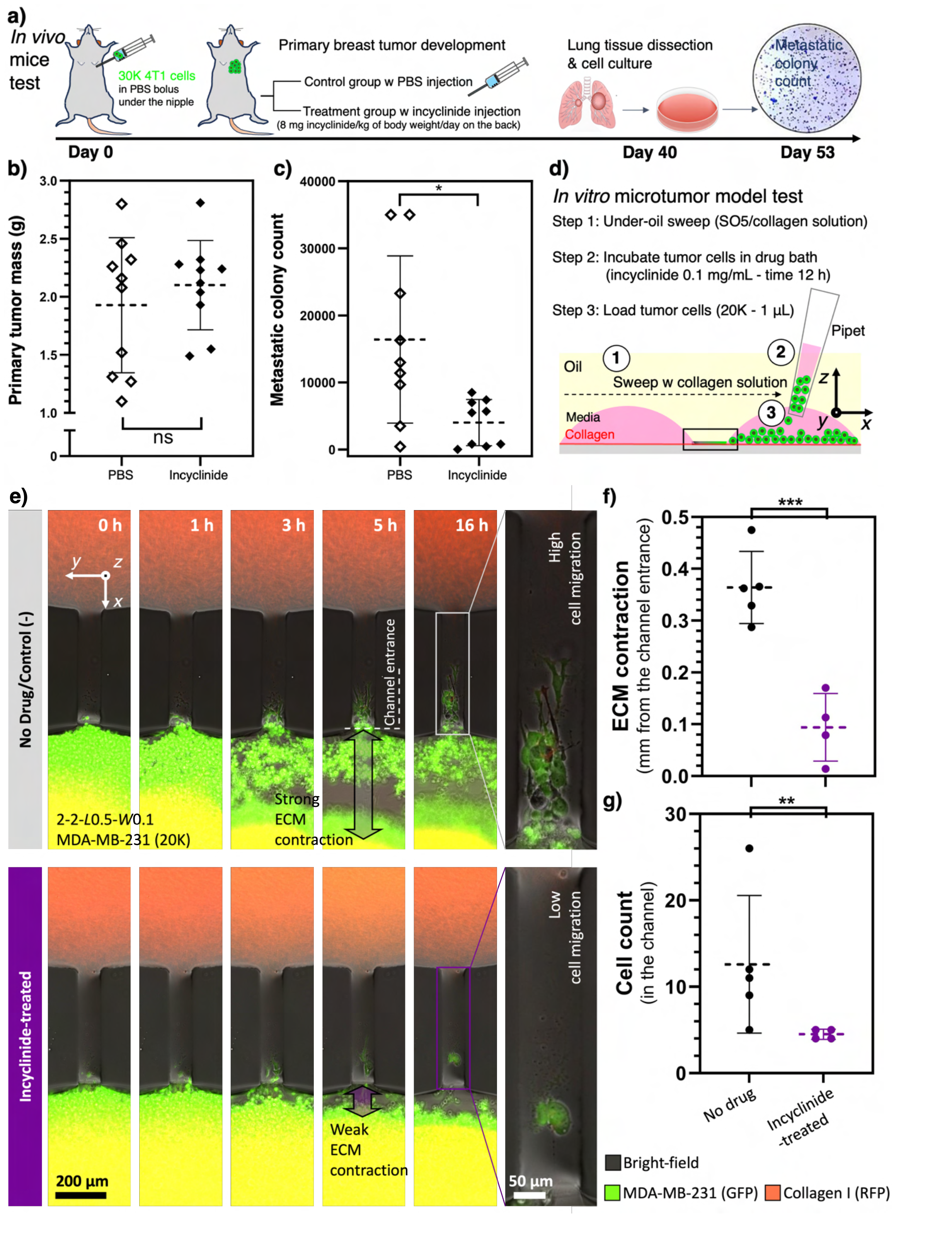
Anti-metastasis drug testing. (a) A schematic illustrates the workflow of *in vivo* mice experiments (see Experimental Section). (b) Primary tumor mass shows no significant differences between the PBS [i.e., control (-)] and incyclinide-treated mice groups. (c) The number of metastatic colonies from the incyclinide-treated group is significantly lower than that of the PBS group. (d) A schematic demonstrating the workflow of UOMS microtumor model test with incyclinide treatment (see Experimental Section). (e) The representative composite microscopic images from the UOMS microtumor model (2-2-*L*0.5-*W*0.1) test with and without incyclinide treatment (Movie S3). (f), (g) ECM contraction and cell count analyses of (e). On UOMS microtumor model, tumor cells treated with incyclinide displayed reduced migration and ECM contraction. Error bars are mean ± S.D. from ≥3 replicates. ^*^*P* ≤ 0.05, ^**^*P* ≤ 0.01, ^***^*P* ≤ 0.001, and ^****^*P* ≤ 0.0001. “ns” represents “not significant”.

## Conclusion

*In vitro* microtumor models^[1]^ continue to play an important role in cancer pathology studies and clinical applications, e.g., patient stratification, drug test and treatment screening. While a decent cluster of *in vitro* microtumor models has been reported, most of these models were developed based on closed-system microfluidics.^[43]^ The closed-system configuration is not well aligned with the open system standard in biology and biomedicine, adding adoption and implementation barriers to the end users (e.g., biologists, oncologists) where the most impact is expected to see from using these models.

In this work, we aim to leverage the ELR-empowered UOMS^[12,21–28]^ to establish an open microfluidic *in vitro* microtumor model with controlled, self-organized ECM microenvironments for studying tumor cell-ECM interactions and pre-screening of anti-metastasis strategies. Our design principle is driven by combined top down and bottom up for synergized controllability and efficiency. The operators are only required to design the surface patterns top down. Defined by the Laplace pressure-driven lateral flow dynamics, a layer of ECM can be readily introduced into an array of open microchannels [e.g., 24 units per standard chambered coverglass (6 cm × 2.4 cm in footprint), max. 4 pieces of chambered coverglass per run of imaging] by a fast sweep^[24,25]^ (in a minute) with either homogeneous or heterogeneous compositions and structures. The following polymerization and self-organization of the ECM microfibers occur spontaneously on the UOMS microtumor device bottom up. The ELR physics and absolute antifouling property^[21]^ make the device fabrication and sample operation/on-device manipulation error-proof, which significantly lowers the adoption and implementation barriers. Cells and reagents (e.g., fresh/conditioned culture media, drug solution, pathogens, etc.) can be easily added to or collected from the units of interest on a device with minimized system disturbance.

This UOMS microtumor model can be further integrated with other UOMS-based technologies including autonomously regulated oxygen microenvironment (AROM),^[12]^ microliter whole blood assay (μ-Blood),^[44,45]^ microbial communities,^[26,46]^ ELR-antimicrobial susceptibility testing (AST),^[27]^ micro total analysis system (μTAS),^[22]^ and label-free spectroscopies (e.g., multiphoton,^[26]^ Raman,^[28]^ and IR^[47]^). Thanks to its natural alignment with the open system standard, the UOMS microtumor model is compatible with most the automated liquid handling tools (e.g., robotic pipetting/microtiter plate preparation systems) in biology and clinical laboratories. Integrated with lab automation will further increase the efficiency, consistency, and throughput of the UOMS microtumor model.^[23]^

## Experimental Section

### Preparation of the ELR-empowered UOMS devices

Detailed protocol was described in our previous publications.^[24,25]^ Briefly, the process includes i) surface modification, ii) surface patterning, and iii) under-oil sample loading. Surface modification introduces a monolayer of covalently bonded PDMS-silane (1,3-dichlorotetramethylsiloxane, Gelest, SID3372.0) molecules by vacuum chemical vapor deposition (CVD) (Bel-Art F420220000, Thermo Fisher Scientific, 08-594-16B) at RT. In this work, we used standard chambered coverglass [Nunc Lab-Tek-II, 2 well (155379), #1.5 borosilicate glass bottom, 0.13 to 0.17 mm thick; Thermo Fisher Scientific] to prepare the devices. The following surface patterning step is to transfer a designated pattern from a PDMS stamp (or mask) to the PDMS-silane grafted surface by selective oxygen plasma treatment (100 W/1 min) (Diener Electronic Femto, Plasma Surface Technology). Finally, oil (silicone oil, 5 cSt, 317667, Sigma-Aldrich) is added to the device and cellular/molecular samples can be loaded under-oil by under-oil sweep distribution or regular pipetting.

### Under-oil microchannels with ECM coating

We prepared the under-oil ECM microchannels by under-oil sweep distribution. Collagen gel (Rat Tail, Type I, CLS354249, Corning) was prepared with 1:1 volume ratio of the original collagen stock (∼10 mg/mL) to 2% (wt.) 2-[4-(2-hydroxyethyl)piperazin-1-yl]ethanesulfonic acid (HEPES) 2×phosphate-buffered saline (PBS) solution, followed by another 1:1 volume ratio dilution with the target cell culture media. The final collagen solution was supplemented with 5% (vol.) fluorescent collagen (RFP or GFP) and lastly adjusted to pH 7.4 with 0.5 M sodium hydroxide (NaOH) endotoxin-free aqueous solution. All these operations were done on ice to prevent pre-polymerization. After sweeping, the ECM microchannel devices were left at RT (∼21 °C) for 1 h to allow a complete polymerization.

### Cell culture and passage

MDA-MB-231 cells (GFP labeled) were cultured in Dulbecco’s modified Eagle’s medium (DMEM) (11960051, Thermo Fisher Scientific) + 10% fetal bovine serum (FBS) (10437010, Thermo Fisher Scientific) and 1% Penicillin-Streptomycin (15140122, Thermo Fisher Scientific) in a standard carbon dioxide (CO_2_) incubator [at 37 °C with 18.6% O_2_, 5% CO_2_, 95% relative humidity (RH)] in a T75 tissue culture flask. When reaching ∼50% confluency, the cells were first washed with 1×PBS then trypsinized with 0.05% Trypsin (25200056, Thermo Fisher Scientific). The released cells were counted and resuspended to 20K cells/μL and then diluted down to 5K cells/μL stocks. 1 μL of the cell suspension was loaded to the cell seeding spot by regular pipetting through the oil overlay. The UOMS microtumor model devices were kept in a standard CO2 incubator before and during the imaging gaps.

### In vitro incyclinide treatment and cell migration

40 mg of incyclinide (TA9H94533373, Sigma-Aldrich) was dissolved in 1 mL dimethyl sulfoxide (DMSO) to reach 40 μg/μL. 1 μL of the incyclinide solution was added to each 10 mL of culture media. After cell passage, the cells were incubated with the incyclinide-supplemented culture media for overnight. The incyclinide-treated cells were first washed with 1×PBS then trypsinized with 0.05% Trypsin. The released cells were counted and resuspended to 20K cells/μL and then diluted down to 5K cells/μL stocks. 1 μL of the cell suspension was loaded to the cell seeding spot by regular pipetting through the oil overlay.

### In vivo incyclinide treatment and tumor metastasis

The mice model experiments were performed in the Liu Lab (Liberty University). 20 of 8-week-old BALB/c mice (Jackson Laboratory) were inoculated with 4T1 mouse mammary cancer cells at 30K cells per mouse in the fat pad under nipple number 9. 10 days later, when tumors became palpable, 10 mice were treated with incyclinide dissolved in 1×PBS with a pH of 7.8. Incyclinide was administered subcutaneously at 8 µg/Kg daily. Control mice received the save volume of 1×PBS at the same pH. Mice were sacrificed 40 days after inoculation (30 days of treatment). The organs of the mice were weighed to calculate organ index (defined as the ratio of individual organ weight to that of the total body weight). Lungs were minced, digested with 1×PBS solution containing collagenase IV (17104019, Thermo Fisher Scientific) at 1 mg/mL and elastase at 6 U/mL (E-240, Goldbio) for 1 h. Cell suspension was filtered using a 70-µm cell strainer (43-50070-51, PluriSelect) and cultured in RPMI 1640 (11875093, Gibco, Thermo Fisher Scientific) supplemented with 10% FBS, 6-thioguanine at 10 µg/mL, and Gibco Antibiotic-Antimycotic (15240062, Thermo Fisher Scientific). Colonies of metastatic cells were stained with crystal violet and counted after 13 days of incubation. The Mann-Whitney U test was used for statistical differences.

### Imaging and time lapse

Bright-field, fluorescence images and videos were recorded on a Nikon Ti Eclipse inverted epifluorescence microscope (Nikon Instruments). The UOMS microtumor model devices were kept at 37 °C, 21% O_2_, 5% CO_2_, 95% RH via an on-stage incubator (Bold Line, Okolab) during imaging. A Zeiss Apotome 3 epifluorescence microscope was used to record the quasi-confocal fluorescence images.

### Image/video analyses

Image/video analyses were performed in Fiji ImageJ. i) Surface area analysis. Regions of interest (ROIs) were drawn manually on the target ECM regions in ImageJ and then measured. ii) Orientation analysis. The orientation analysis of the ECM microfibers was performed in ImageJ using the OrientationJ plugin. The color survey images were generated with “OrientationJ Analysis”. The coherency was measured with “OrientationJ Measure”. iii) Cell morphology analysis. The circularity of cells was analyzed in ImageJ using the “Analyze → Analyze Particles…” function. iv) Cell counting and speed tracking analysis. Manual counting was performed in ImageJ with the timelapse videos recorded on the Nikon microscope by counting each individual cell that passes into the channel. For speed tracking, images in a timelapse were transferred into ImageJ, where cell tracking was performed manually using the MTrackJ plugin. Once MTrackJ is selected, the “Add” option allows tracking of each individual migrating cell. Once the manual tracking is done for each migrating cell, measurements can be taken automatically. Measurements were created through the MTrackJ measurements option. All data graphs were prepared in GraphPad Prism 10.1.2. Videos were edited and compiled in VSDC video editor.

### Statistical analysis

Raw data was directly used in statistical analysis with no data excluded. Data was averaged from at least 3 replicates (unless otherwise stated) and present as mean ± standard deviation (S.D.) if applicable. The statistical significance was specified in the figure captions. All statistical analyses were performed using GraphPad Prism 10.1.2.

## Acknowledgements

Experimental work was performed in the MMB Lab facility at the University of Wisconsin-Madison. We thank Dr. David J. Beebe for offering suggestions. The mice experiments were supported by Dr. Yingguang Liu and performed in the Liu Lab at the College of Osteopathic Medicine, Liberty University. We thank Dr. Adeel Ahmed (the MMB Lab) for the discussion on the recent progress in microfluidics-based ECM alignment techniques. Funding: This work was supported by Liberty University Center for Research and Scholarship. Author contribution: C.L. conceived the project and designed the research. C.L. directed the under-oil microtumor model development and C.L., J.L., Z.A., M.A.F., and B.R. performed data collection. J.L., A.K., S.N. and Y.L. designed and conducted the *in vivo* mice experiments. C.L., J.L. and N.W.H. performed data analysis and visualization. C.L. supervised the project. C.L. and J.L. wrote the manuscript and all authors revised it.

## Data Availability Statement

All study data are included in the article and/or supporting information. The data that support the findings of this study are available from the corresponding authors upon reasonable request.

