## Supporting Information for "Combining Top-Down and Bottom-Up: An Open Microfluidic Microtumor Model for Investigating Tumor Cell-ECM Interaction and Anti-Metastasis"

**This SI PDF file includes:**

Figures S1 to S4
Table S1
Legends for Movies S1 to S3

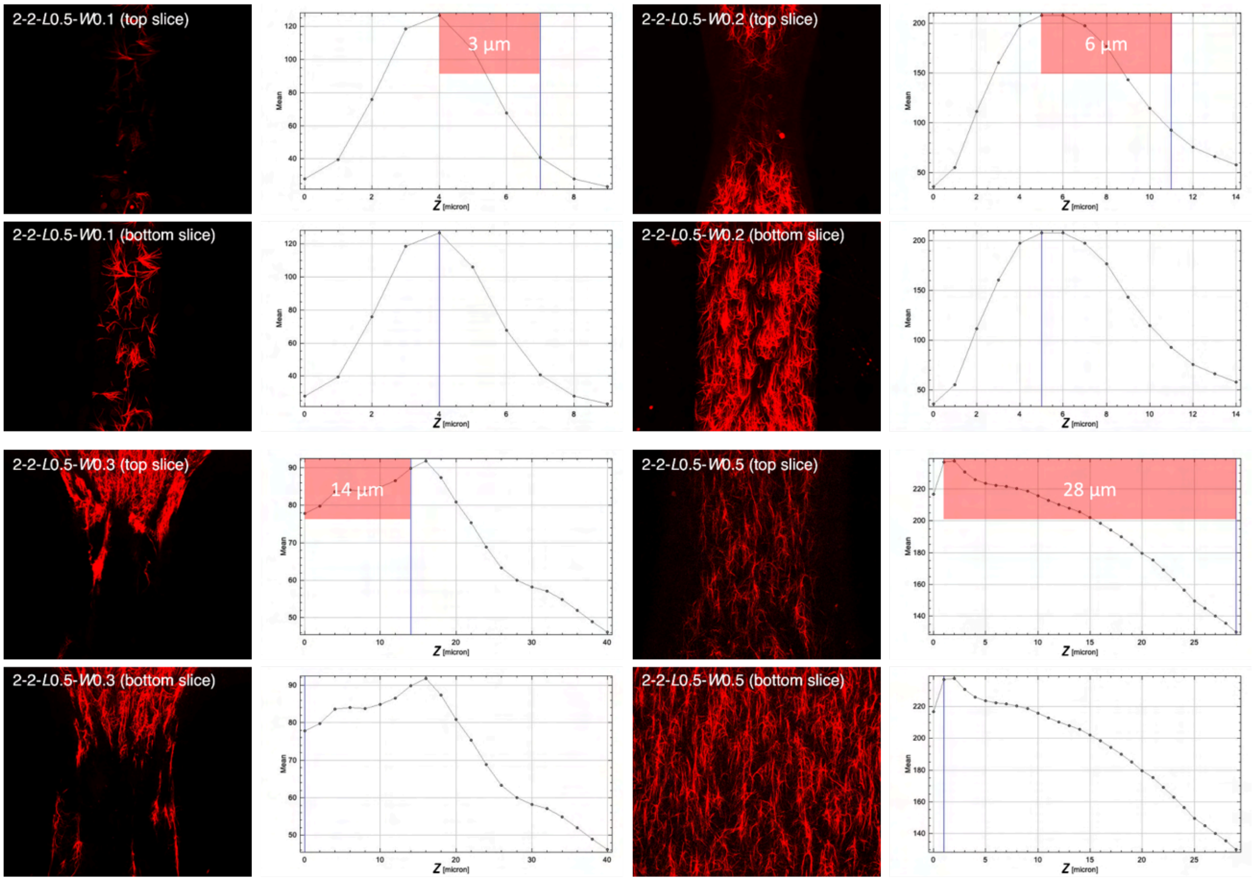


***Figure S1.*** ECM layer thickness measurements using Zeiss Apotome (i.e., quasi-confocal) microscopy. The ECM layer was scanned in z-stack at a step of 2 µm from bottom to top. The bottom slice and top slice of the ECM layer in a microchannel were identified as the first image with the most microfibers appearing in the field of view and the first image the microfibers at the middle of the channels disappearing in the field of view, respectively. The corresponding distance in z between the bottom slice and top slice on the intensity (mean)-z plots is recorded as the ECM layer thickness. At least 3 replicate channels were measured and averaged for each channel dimension (Figure 2b, red line).

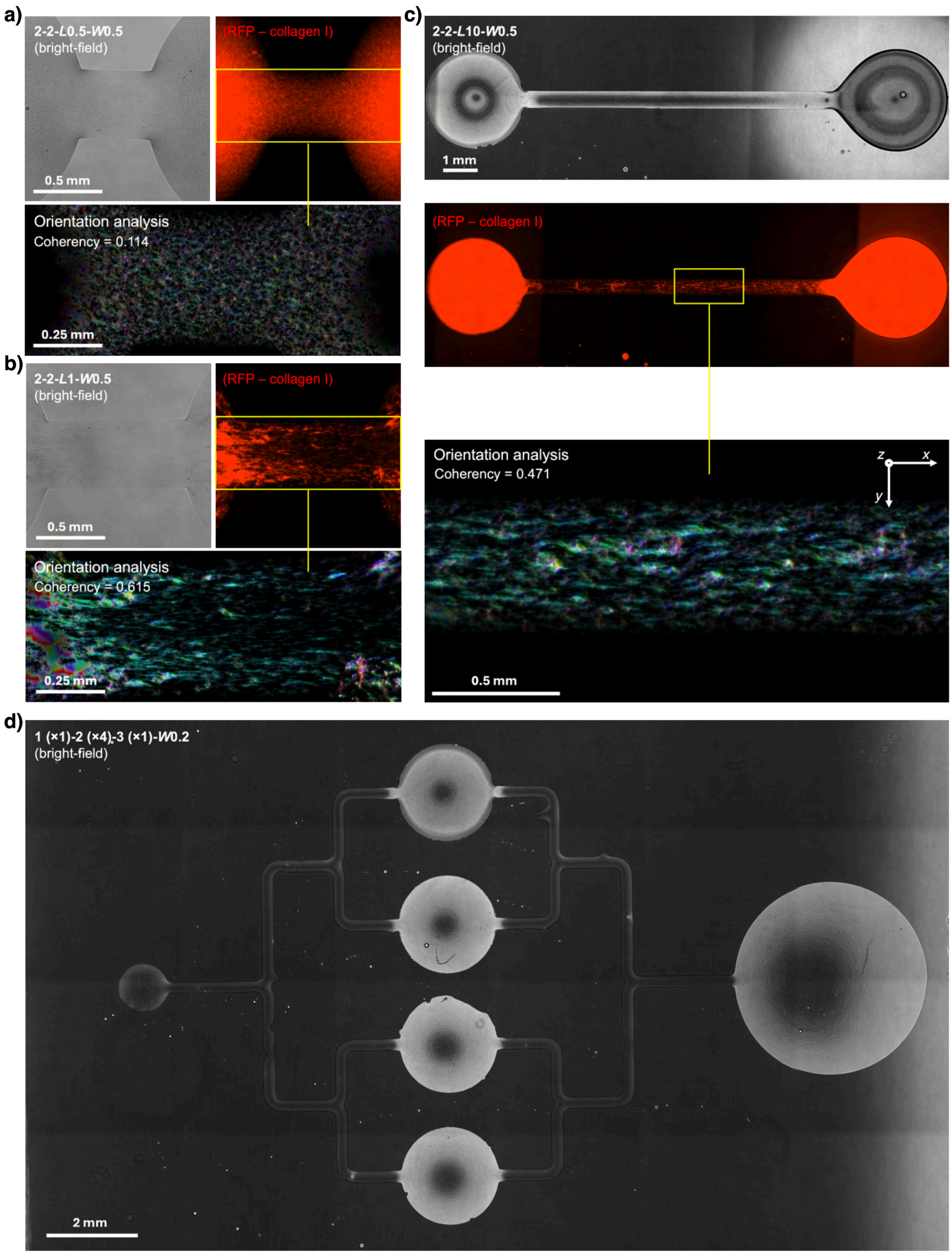

a)
2-2-L0.5-W0.5
(bright-field)
0.5 mm
(RFP – collagen I)
Orientation analysis
Coherency = 0.114
0.25 mm
b)
2-2-L1-W0.5
(bright-field)
0.5 mm
(RFP – collagen I)
Orientation analysis
Coherency = 0.615
0.25 mm
c)
2-2-L10-W0.5
(bright-field)
1 mm
(RFP – collagen I)
Orientation analysis
Coherency = 0.471
z
x
y
0.5 mm
d)
1 (×1)-2 (×4)-3 (×1)-W0.2
(bright-field)
2 mm

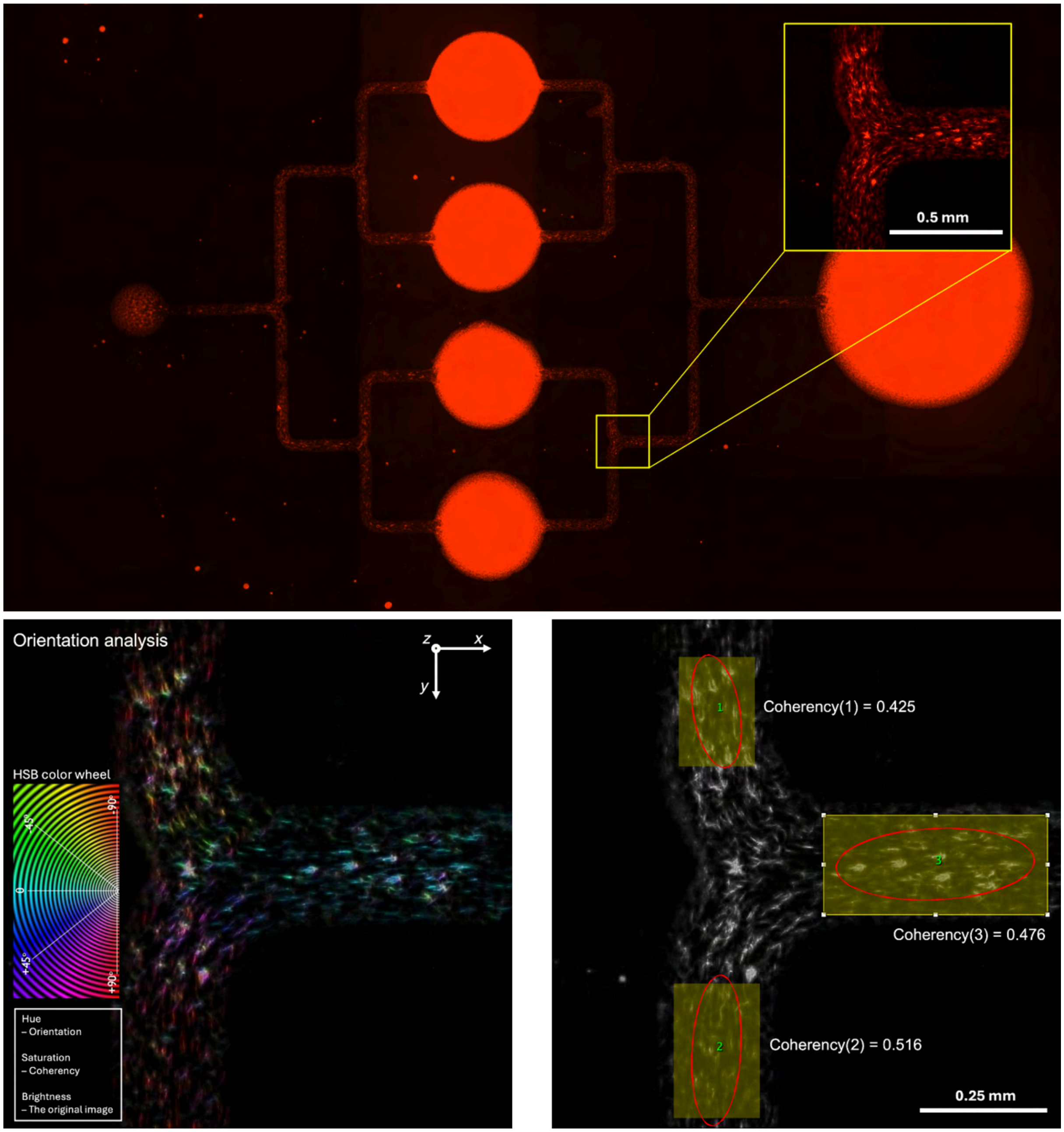
0.5 mm
Orientation analysis
z
x
y
HSB color wheel
-45°
-90°
0
+45°
+90°
Hue
– Orientation
Saturation
– Coherency
Brightness
– The original image
Coherency(1) = 0.425
Coherency(3) = 0.476
Coherency(2) = 0.516
0.25 mm

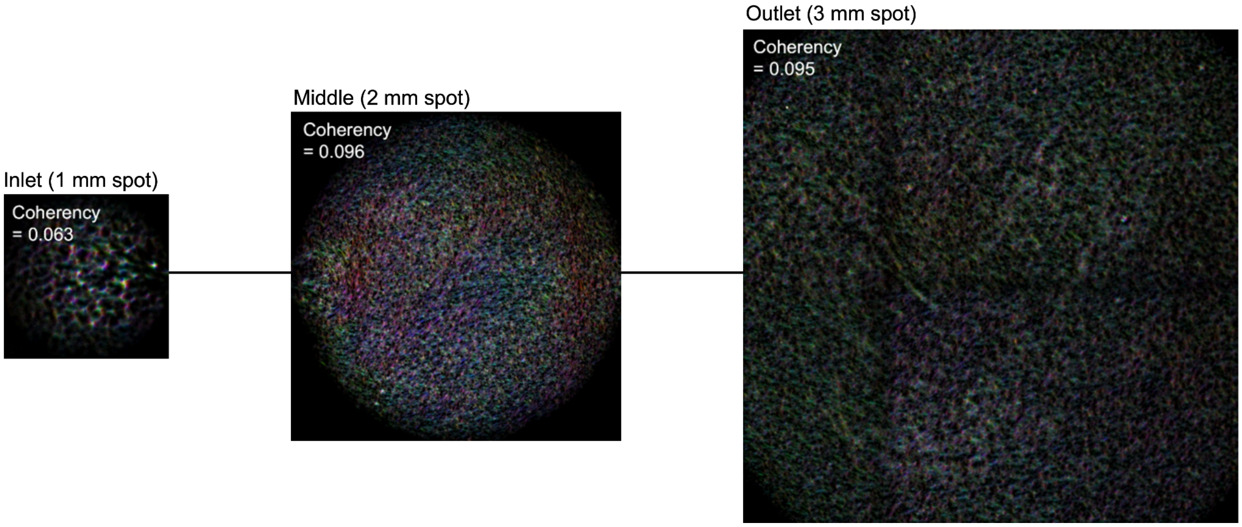


***Figure S2.*** Influence of channel length on collagen microfiber alignment.(a), (b), and (c) Microchannels with 0.5 mm in channel width and varying channel lengths at 0.5 mm, 1 mm, and 10 mm. The ECM microfiber alignment in the microchannels were measured (see Experimental Section) and compared. (d) A complex channel with 1 of 1 mm inlet spot, 4 of 2 mm middle spots, and 1 of 3 mm outlet spot. The ECM microfiber alignment in the microchannels and on the spots were measured and compared.

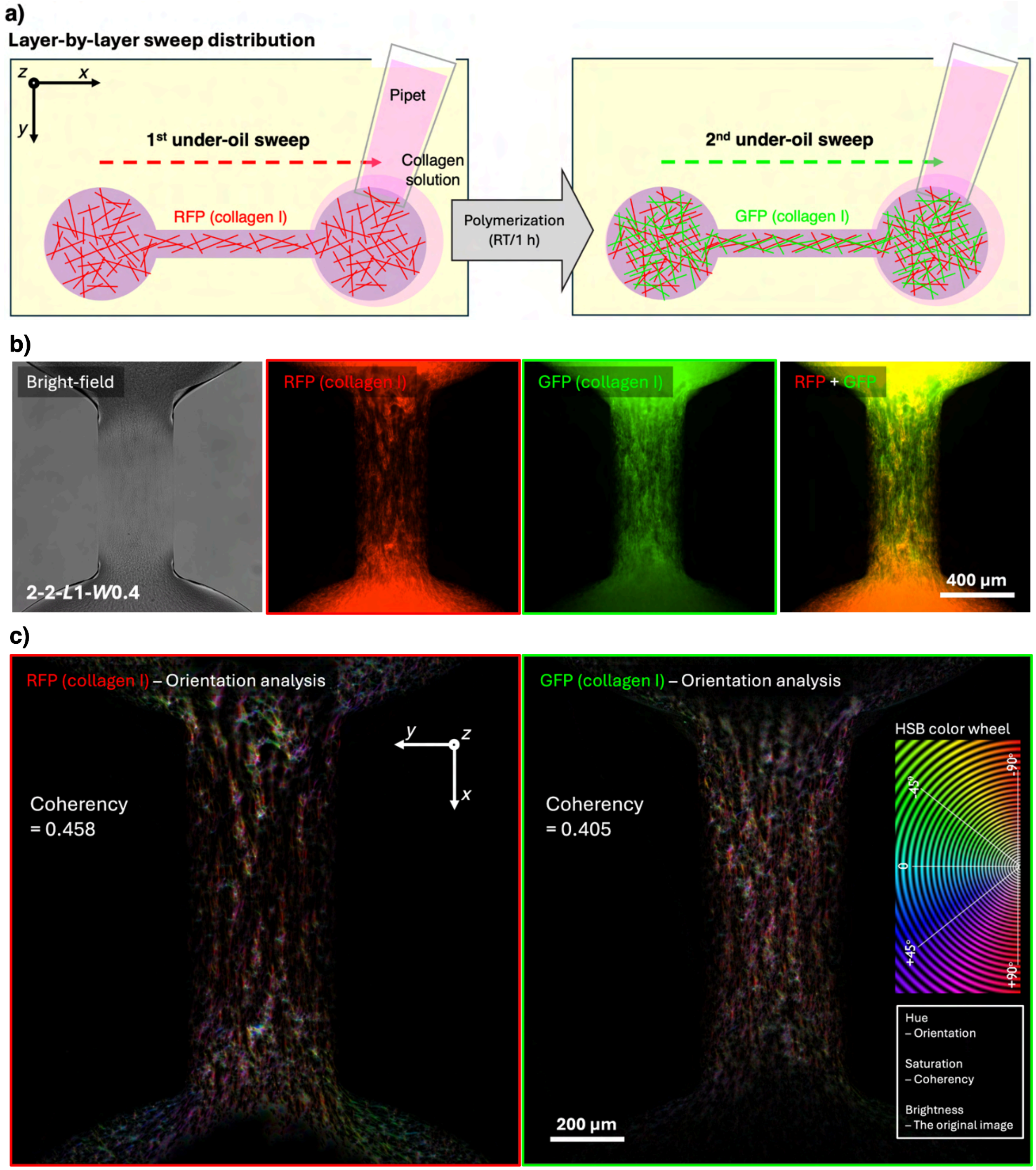


***Figure S3.*** Layer-by-layer sweep distribution of collagen (Type I) in a microchannel. (a) A schematic showing the layer-by-layer sweep distribution. (b) The microscopic images of bright field and the fluorescent (RFP and GFP) channels. (c) The orientation analysis (see Experimental Section) of the ECM layers. The results showed that the second sweep successfully applied another layer of ECM on the first layer with similar microfiber alignment but different compositions.

a)

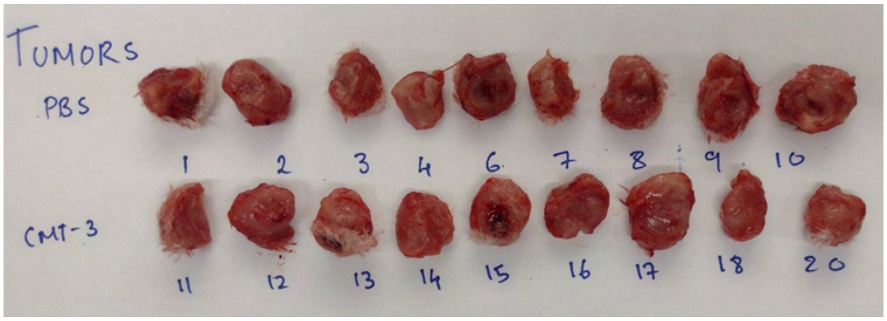
TUMORS
PBS
1
2
3
4
6
7
8
9
10
CMT-3
11
12
13
14
15
16
17
18
20

b)

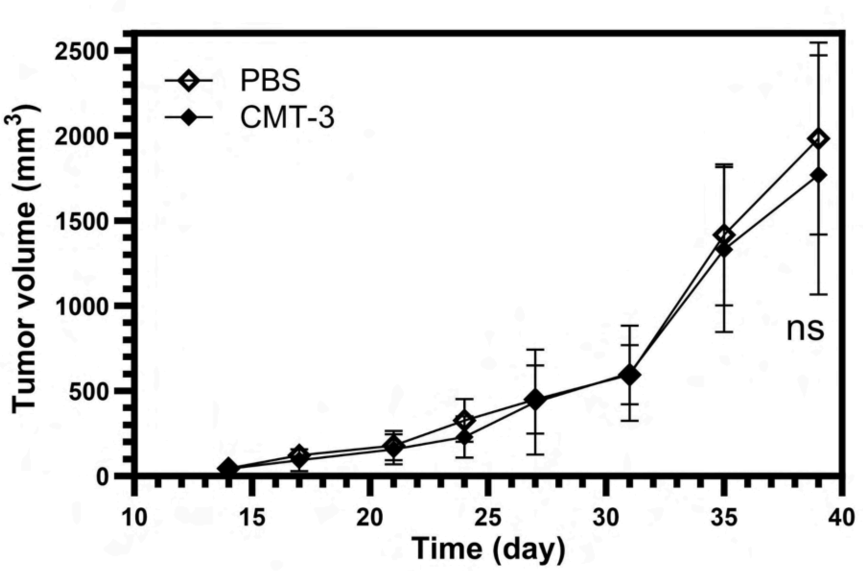
PBS
CMT-3
Tumor volume (mm$^3$)
Time (day)
ns
0
500
1000
1500
2000
2500
10
15
20
25
30
35
40

c)

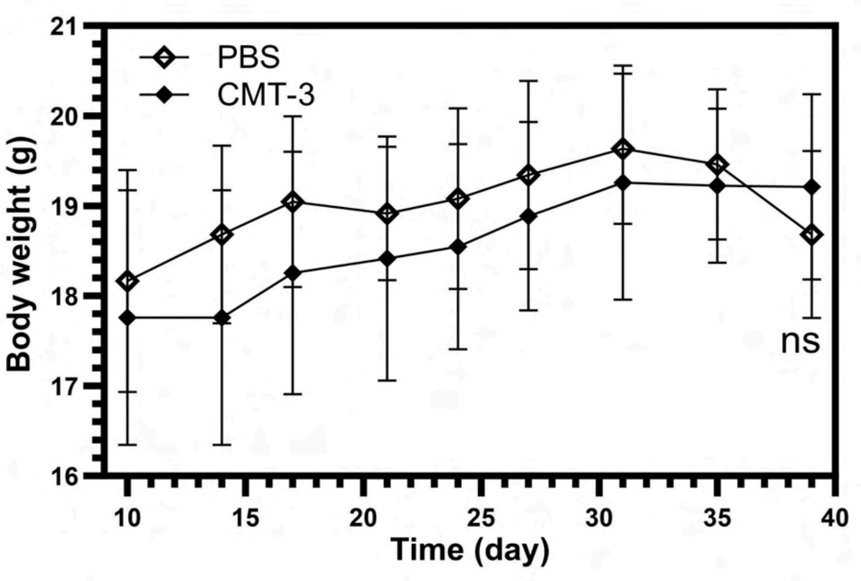
PBS
CMT-3
Body weight (g)
Time (day)
ns
16
17
18
19
20
21
10
15
20
25
30
35
40

**d)**

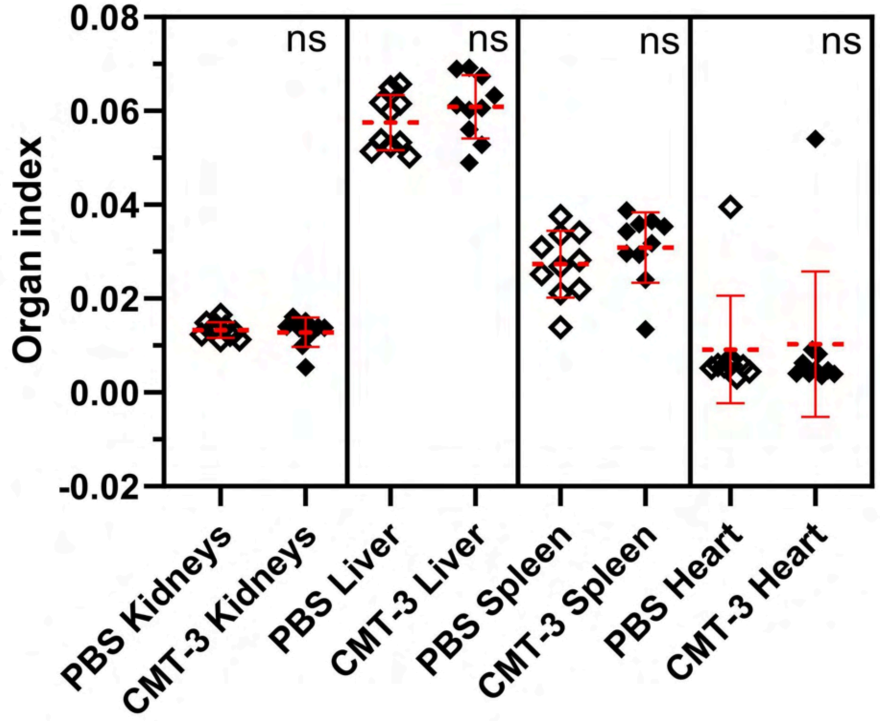


***Figure S4.*** *In vivo* mice experiment results. (a) Primary breast tumor specimen from mice in incyclinide-treated and PBS control groups harvested on Day40. No significant difference was observed grossly with the primary tumors between the treatment and control groups. (b) Tumor volume measurement from Day14 to Day40 showed no significant difference between the treatment and groups. (c) There was an initial increase in average body weight in both the treatment and control groups due to the enlarging tumor mass. The plateau towards the end of the timeline is attributed to tumor burden. (d) Organ index (see definition in Experimental Section) of different organs from the mice test. The results showed no significant difference between the treatment and control groups, indicating no apparent drug-related organ damage associated with incyclinide treatment.

***Table S1.*** Fluid parameters right after sweep.

| I - Right after sweep (see Figure 2a) | | | | | |
|---|---|---|---|---|---|
| *W* (mm) | *H* (*H*/*W* = 1/13) (μm) | *L* (mm) | *R* (mm)* | Cross-section ($\mu m^2$) | $V_{channel}$ (pL) |
| 0.1 | 7.7 | 0.5 | 0.166 | 515.2 | 257.6 |
| 0.2 | 15.4 | 0.5 | 0.333 | 2061.0 | 1030.5 |
| 0.3 | 23.1 | 0.5 | 0.499 | 4637.1 | 2318.6 |
| 0.5 | 38.5 | 0.5 | 0.831 | 12881.0 | 6440.5 |
| 2.0 ($D_{spot}$) | 153.8 | N/A | 3.328 | N/A | 243000 |

*For the microchannels, *R* in the table is the radius of curvature in the channel width direction, i.e., $R_{channel\text{-}width}$.

***Movie S1.*** ECM remodeling and alignment in the microchannel driven by tumor cells.

***Movie S2.*** (a) Individual cell migration at low seeding density. (b) Cell cluster slingshot at high seeding density. (c) ECM contraction and remodeling.

***Movie S3.*** Comparison between (a) no drug and (b) incyclinide treatment on ECM contraction and tumor cell migration.
